# Structure-based modelling and dynamics of MurM, a *Streptococcus pneumoniae* penicillin resistance determinant that functions at the cytoplasmic membrane interface

**DOI:** 10.1101/2020.06.18.158840

**Authors:** Anna York, Adrian. J. Lloyd, Charo I. del Genio, Jonathan Shearer, Karen. J. Hinxman, Konstantin Fritz, Vilmos Fulop, Syma Khalid, Christopher. G. Dowson, David. I. Roper

## Abstract

MurM is an aminoacyl-tRNA dependant ligase that aminoacylates the Lipid II peptidoglycan precursor, in the human pathogen *Streptococcus pneumoniae*. MurM is required for the generation of branched peptidoglycan precursors enabling indirect cross-links in the peptidoglycan and is found to be essential for penicillin resistance. In this study we have solved the X-ray crystal structure of *Staphylococcus aureus* FemX, an isofunctional homologue of MurM, and used this as a template to generate a homology model of MurM. Using this model, we perform molecular docking and molecular dynamics to examine the interaction of the protein with the phospholipid bilayer and the membrane embedded Lipid II substrate of MurM. Our model suggests that MurM is associated with the major membrane phospholipid cardiolipin, and we confirm this with experimental evidence that the activity of MurM is enhanced by this phospholipid and inhibited by its direct precursor phosphatidylglycerol. This suggests that the spatial association of pneumococcal membrane phospholipids and their impact on MurM activity may be a critical to the final architecture of the peptidoglycan and the expression of clinically relevant penicillin resistance in this pathogen.

## Introduction

The peptidoglycan (PG) of the bacterial cell wall is a polymer consisting of alternating *β*-1,4 linked *N*-acetyl glucosamine (Glc*N*Ac) and *N*-acetyl muramic acid (Mur*N*Ac) residues. Appended to the Mur*N*Ac sugar is a pentapeptide stem that can be cross-linked directly or indirectly to form a rigid mesh-like structure (Bugg et al., 2011). PG biosynthesis begins with the cytoplasmic formation of a Park nucleotide, which is subsequently converted into a lipid-linked PG precursor known as Lipid II. Lipid II is then transported across the membrane, where it is polymerised and cross-linked by the penicillin-binding proteins (PBPs) (Figure 1). PG is an essential component of the cell wall, involved in cell growth and division, maintaining structural integrity, and resisting high osmotic pressures. Inhibition of cell wall biosynthesis is a key mechanism for many antibiotics, including β-lactams, glycopeptides and amino acid analogues (Schneider and Sahl, 2010).

**Figure 1:**
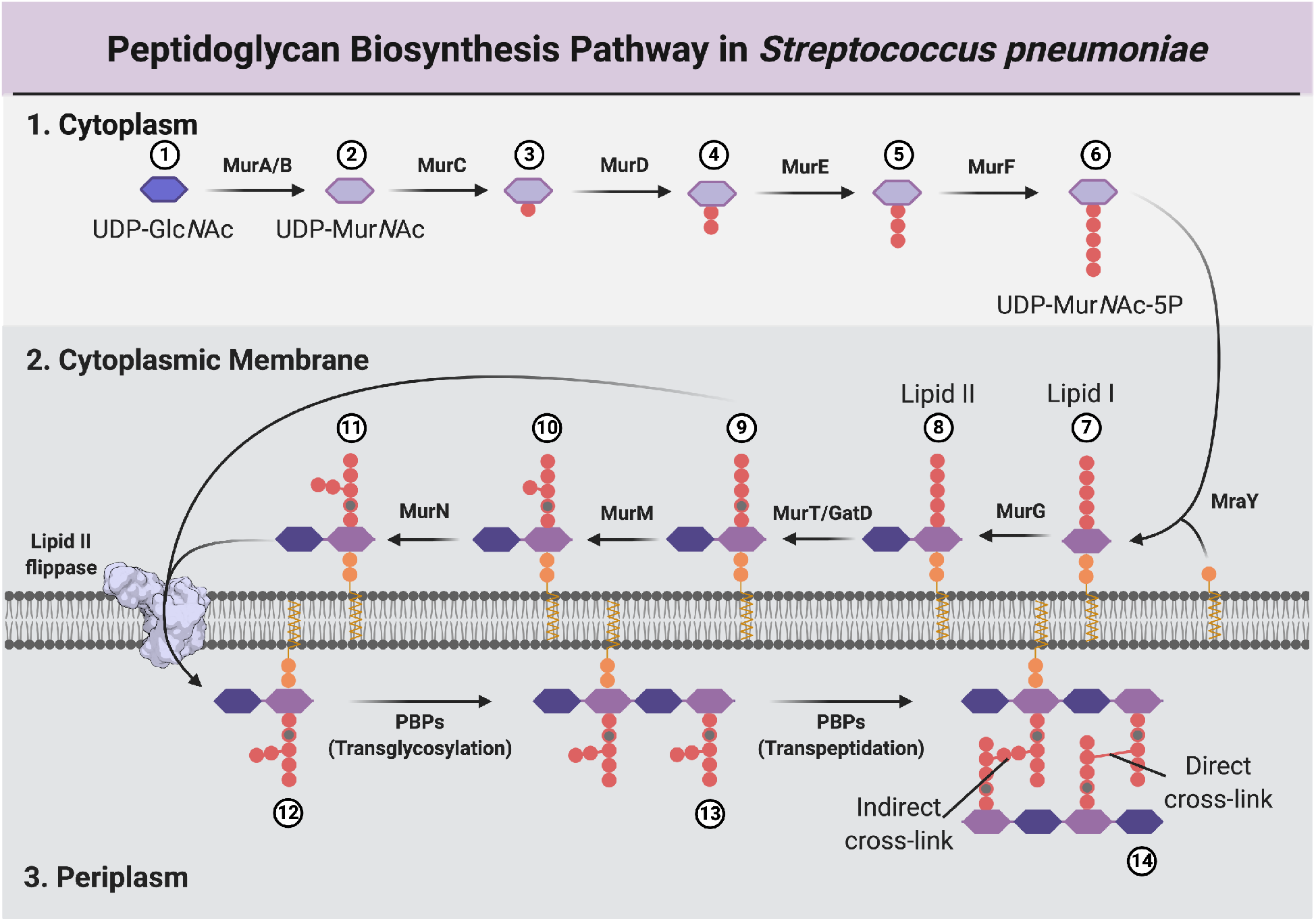
Diagram illustrating stages of the PG biosynthesis pathway. 1) The cytoplasmic stage is characterised by the formation of UDP-Mur*N*Ac-pentapeptide (UDP-Mur*N*Ac-5P) by the Mur ligases. The pentapeptide (5P) stem peptide usually comprises L-Ala-γ-D-Glu-L-Lys-D-Ala-D-Ala in Gram-positive organisms. 2) At the internal face of the cytoplasmic membrane MraY catalyses the addition of UDP-Mur*N*Ac-5P to undecaprenyl-pyrophosphate forming Lipid I, which is then converted to Lipid II by MurG. In *S. pneumoniae*, the second position D-glutamate is α-amidated to D-iso-glutamine (iGln) by the MurT/GatD complex, and in some cases a dipeptide branch of either L-Ser/L-Ala or L-Ala/L-Ala may be appended at the e-amino group of the third position lysine by MurM and MurN, respectively. The exact order of the cytoplasmic membrane steps remains uncertain, but for clarity, in this figure, they appear in a linear format, with conversion to Lipid II occurring before peptide stem modifications, and amidation occurring before branching. Lipid II is translocated across the membrane by MurJ. 3) At the external face of the cytoplasmic membrane, PBPs form glycan chains by transglycosylation (TG), with the concomitant release of undecaprenyl-pyrophosphate, and form either direct or indirect cross-links throughout the PG layer via transpeptidation (TP). Nucleotide sugars UDP-Glc*N*Ac and UDP-Mur*N*Ac and the sugars *GlcN*Ac and Mur*N*Ac are signified by blue, violet, dark blue and purple elongated hexagons respectively. Figure created with BioRender.com.

In *S. pneumoniae* and other Gram-positive bacteria, the glutamate at the second position of the Lipid II pentapeptide is α-amidated to *iso*-glutamine by the essential GatT/MurD complex (Figueiredo et al., 2012, Münch et al., 2012, Zapun et al., 2013, Morlot et al., 2018). In addition, branched Lipid II, capable of generating indirect crosslinks, can be formed by the non-essential MurM and MurN proteins. MurM and MurN are responsible for the sequential addition of amino acids to the third-position lysine of the pentapeptide stem (Filipe et al., 2000). MurM can append either L-serine or L-alanine at the first position of the dipeptide bridge, whilst MurN extends this modification by addition of an invariable L-alanyl moiety. Branched PG precursors are also found in several other Gram-positive bacterial pathogens, for example the glycyl-tRNA^Gly^ dependent enzymes FemX, A and B are responsible for the addition of a pentaglycyl bridge in *Staphylococcus aureus* (Schneider et al., 2004). In comparison with other Gram positive organisms, the PG of *S. pneumoniae* is highly heterogeneous: the predominant C-terminal amino acid at position 1 of the dipeptide, and the proportion of indirect cross-links throughout the PG, vary significantly between different strains (Severin and Tomasz, 1996, Garcia-Bustos et al., 1987, Garcia-Bustos and Tomasz, 1990). *In vitro* and *in vivo* studies indicate that MurM from the penicillin-resistant strain *S. pneumoniae*(159) preferentially incorporates L-alanine, whilst MurM from a penicillin-sensitive strain *S. pneumoniae* (Pn16) preferentially incorporates L-serine (Lloyd et al., 2008). In addition, penicillin resistant strains demonstrated higher levels of indirect cross-linking in the PG compared to penicillin susceptible isolates, however the overall degree of cross-linking remained constant (Garcia-Bustos and Tomasz, 1990).

Resistance to β-lactam antibiotics in *S. pneumoniae* is characterised by extensive interspecies recombination of PBP transpeptidase domains, which results in a mosaic PBP active site with a lower β-lactam binding affinity (Smith et al., 1991). This mechanism of resistance contrasts with that of many other bacteria that have acquired genes for β-lactamase enzymes which inactivate the antibiotic before it binds to and inhibits the PBPs. Interestingly, deletion of the *murM* gene in *S. pneumoniae* eliminates indirect cross-links from the PG and results in a complete loss of penicillin resistance (Filipe et al., 2001). It has been proposed that the changes to the PBP active site which prevent β-lactam binding, may also alter the Lipid II substrate specificity such that the PBPs bind branched Lipid II more tightly than unbranched Lipid II. MurM is therefore necessary, but not sufficient for resistance in clinical strains of *S. pneumoniae*, making it an interesting target for the development of new inhibitors of antimicrobial resistance (Filipe and Tomasz, 2000).

The cytoplasmic membrane of *S. pneumoniae* contains two phospholipids, phosphatidylglycerol and cardiolipin (Trombe et al., 1979, Pesakhov et al., 2007), where cardiolipin synthase is responsible for generating cardiolipin from two molecules of phosphatidylglycerol (Schlame, 2008). The proportion of cardiolipin and phosphatidylglycerol, as a percentage of the overall membrane lipids, varies in *S. pneumoniae* between anaerobic and aerobic growth conditions. Cardiolipin was found to decrease from 15.3 % to 8.3 %, whilst phosphatidylglycerol increased from 12.7 % to 16.3 % in anaerobic conditions compared to aerobic conditions (Pesakhov et al., 2007). The peptidoglycan precursor, Lipid II is tethered to the cell membrane by virtue of its C55 Lipid II tail. MurM then transfers a single amino acyl moiety (Seryl or Alanyl) from aminoacyl-tRNA to the third position lysine of the cytoplasmically oriented Lipid II pentapeptide stem.

Previously, MurM inhibitors have been identified; however, none have shown growth inhibition of effect on penicillin MIC, indicating that these compounds cannot effectively cross the cytoplasmic membrane of *S. pneumoniae* (Cressina et al., 2007, 2009). MurM has thus far resisted extensive crystallization in our laboratory, and consequently, its X-ray-solved structure is not available. However, in a related study we were able to solve the X-ray structure of the isofunctional homologue of MurM from *S. aureus* (FemX), which we have used here as a template for homology modelling of MurM. Using this MurM homology model we have successfully identified the Lipid II binding site, and used molecular dynamics (MD) simulations to investigate interactions between MurM and both membrane phospholipids and its Lipid II substrate (Witzke et al., 2016). We subsequently, studied the effects of these membrane embedded phospholipids, on the enzymatic activity of MurM *in vitro*, corroborating our *in silico* analysis. These studies provide new insights into the structure and activity of MurM, providing a link between phospholipid membrane composition and peptidoglycan architecture. This may be useful for the development of novel chemical probes for these proteins, and have important implications for future studies on penicillin resistance mechanisms in *S. pneumoniae*.

## Materials and Methods

### Cloning, overexpression and purification of S. aureus FemX

The *S. aureus* Mu50 FemX gene was amplified from chromosomal DNA using oligonucleotides FemX forward: TTTGCGGGTGGTCTCCCATGGAAAAGATGCATATCACTAATCAGG and FemX Reverse: TTTGCGCTCGAGGCCCTGAAAAT-ACAGGTTTTCTTTTCGTTTTAATTTACGAGATATTTTAATTTTAGC. The resulting PCR fragment was cleaved with BsaI and XhoI and was cloned into pET28 between the NcoI and XhoI restriction sites to create pET28::FemX, containing a Tobacco Etch Virus (TEV) protease cleavable carboxyl terminal hexa-histidine tag. *Escherichia coli* BL21 Star (DE3) and *E. coli* B834 (DE3) which had previously been transformed with the pACYC based plasmid (pRARE2) which supplies seven rare tRNAs to support expression of genes in *E. coli* were transformed with pET28::FemX. The transformed *E. coli* BL21 Star (DE3) pRARE2 and *E. coli* B834 (DE3) pRARE2 were used to inoculate Luria broth media and M9 media supplemented with all 19 L-amino acids and 40 mM L-selenomthionine in place in L-methionine respectively (Doublié, 1997). Transformants were cultured at 37 °C at 180 rpm in a shaking incubator until an optical density at 600 nm (OD_600nm_) of 0.4-0.6 was reached. Protein expression was induced by addition of 1 mM isopropyl β-D-1-thiogalactopyranoside, concurrent with a reduction in temperature to 25 °C for 4 hours. Cells were harvested by centrifugation at 6,000 *xg* for 15 minutes and pellets were stored at −20 °C until required. Cell pellets containing 4-6 g of cells were resuspended in 20 mL of 50 mM sodium phosphate pH 7.0 and 1M NaCl to which, one tablet of Pierce EDTA free Protease Inhibitor Tablet was added along with 2.5 mg.mL^-1^ lysozyme. The cell suspension was incubated with slow rotation for 30 minutes at 4°C before the disruption by sonication using a Bandelin Sonopuls device with 3 x 30 second bursts at 70 % power. A crude extract was then prepared by centrifugation at (4 °C) at 50,000 *xg* for 30 minutes. FemX was then purified by immobilised metal affinity chromatography (IMAC) using a 5 mL gravity fed column of cobalt Talon resin equilibrated with 50 mL of 50 mM sodium phosphate pH 7.0, 500 mM NaCl, 10 mM imidazole and 20 % (v/v) glycerol (equilibration buffer). Once the 50,000 *xg* supernatant was loaded onto the column, it was eluted sequentially with 50 mL of equilibration buffer, 30 mL of equilibration buffer with 50 mM imidazole and 30 mL of equilibration buffer with 200 mM imidazole. The last two washes were collected in three 10 mL fractions each. Fractions were analysed by SDS-PAGE and those containing FemX were pooled and concentrated, using a vivaspin 20 centrifugal concentrator column with a 10,000 molecular weight cut off (MWCO), as required. Size exclusion chromatography was used to further purify FemX on a Superdex 75 10/300 column attached to an AKTA purifier system. The column was equilibrated in buffer containing 50 mM sodium phosphate pH 7.0, 500 mM NaCl and 20 % (v/v) glycerol. The histidine tag was then removed from the FemX protein by digestion with histidine-tagged TEV protease at a molar ratio of 100:1 FemX: TEV protease at 4°C overnight. Cleaved and uncleaved protein were separated by a reverse IMAC following the procedure described above.

### Crystallisation and Data collection

FemX was exchanged into 50 mM ethanolamine pH 10.0, 100 mM NaCl and 20 % (v/v) glycerol, concentrated to 15 mg.mL^-1^ using a vivaspin 20 centrifugal concentrator column with a 10,000 MWCO and screened for suitable crystallisation conditions using a honeybee 963 crystallisation robot against JCSG plus, PACT primer and Morpheus crystallisation screens. Crystals obtained from the Morpheus screen were used directly for data collection experiments, although crystallization conditions were further refined to 0.12 M Ethylene Glycols, 0.1 M MES/imidazole pH 6.3 and 28 % (w/v) Ethylene Glycol-PEG 8000. Crystals were frozen directly for X-ray diffraction data experiments on the I04-1 beamline at the Diamond synchrotron (Didcot, UK) using a Pilatus 6M-F detector. Data were processed automatically using Xia2 (Winter, 2010) to 1.62 Å. Molecular replacement was not successful so selenomethionine containing FemX crystals (FemX-SeMet) were produced as described above and the structure was solved by the single anomalous diffraction method. X-ray data from the FemX-SeMet crystal were collected on the I02 beamline at the Diamond synchrotron (Didcot, UK) using a Pilatus 6M detector. All data were indexed, integrated and scaled using the XDS package (Kabsch, 2010). All 10 of expected selenium atoms in the asymmetric unit were located and refined by the SHELX suite (Sheldrick, 2010). These sites were used to obtain preliminary phases. The starting model was built by ARP/wARP (Langer et al., 2008). This model was used to refine the higher resolution data. The structure was refined using iterative cycles of REFMAC (Vagin et al., 2004) and model building/solvent addition with COOT (Emsley et al., 2010).

### S. pneumoniae MurM enzymology

MurM from *S. pneumoniae*(159), also referred to as MurMi59, and its substrates were purified, prepared and assayed as described in Lloyd et al. (2008). MurM was assayed in duplicate in a final volume of 35 *μL* of assay buffer (50 mM 3-(N-morpholino)-propane sulphonic acid, 30 mM KCl, 10 mM MgCl_2_, 1.5 % (w/v) CHAPS adjusted to pH 6.8), 1 mM DTT, 1 mM L-alanine, 10 μM Lipid II-Lys and 24.3 nM MurM. Reactions were initiated by the addition of 0.45 μM [^3^H]-alanyl-tRNA^Ala^ (1000 cpm.pmol^-1^) prepared from *Micrococcus flavus* total tRNA as in Lloyd et al. (2008) and were incubated at 37°C for two minutes, over which time frame, product accumulation was linear with respect to time. Where the impact of cardiolipin or phosphatidylglycerol on MurM activity was assessed, the required amounts of 10 mg/mL stocks of each phospholipid in ethanol or chloroform/methanol (49:1) were dried down in the reaction vials the assays were to be performed in, and solubilised by addition of assay buffer. Reactions were terminated by the addition of 35 μL of ice-cold 6 M pyridinium acetate pH 4.5 and 70 μL ice-cold n-butanol. The incubations were rapidly mixed and centrifuged for 5 minutes at 1°C at 13,000 *xg*, after which time the n-butanol phase was washed with 70 μL of water and then assayed for [^3^H]-Lipid II-L-Ala by liquid scintillation counting. Tritium counts accumulated in control reactions performed without Lipid II-Lys were subtracted from corresponding data acquired in the presence of this substrate. MurM activities in the presence of phospholipid were related to the activity of the enzyme in the absence of phospholipid and plotted as fold activation or percentage inhibition vs phospholipid concentration. The data were then fitted using GraphPad Prism to either of equations 1 or 2 as appropriate:

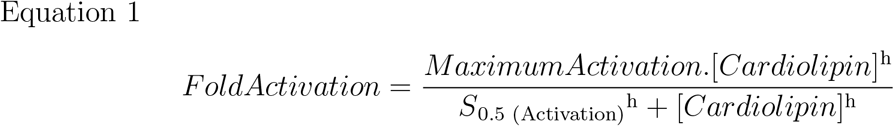

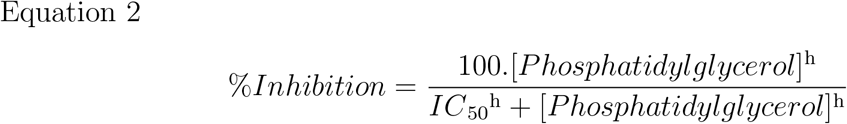

Maximum activation and S_0.5 (Activate)_ (Equation 1) corresponded to the degree of activation at infinite cardiolipin concentration and the cardiolipin concentration required to elicit half maximal activation respectively. IC_50_ (Equation 2) corresponds to the phosphatidylglycerol concentration that elicited half maximal inhibition. For both equations, h denoted the Hill coefficient.

### Homology Modelling of MurM

Due to the natural ability of streptococci to undergo homologous recombination, *S. pneumoniae MurM* genes are highly mosaic, and so, in line with the enzymology studies, the MurM sequence used for homology modelling was that of *S. pneumoniae* MurM_15g_ (Supplementary Material: Figure S1). *S. pneumoniae* MurM, *S. aureus* FemX (PDB ID: 6SNR), *S. aureus* FemA (PDB ID: 1LRZ) and *Weissella viridescens* FemX (PDB ID: 3GKR) were aligned by pairwise sequence alignment using EMBOSS Needle (Madeira et al., 2019) and sequence identity and similarity determined.

The structure of *S. aureus* FemX was used as the template for homology modelling due to its high relatedness with MurM. *S. aureus* FemX and MurM_159_ sequences were aligned, and using MODELLER (Eswar et al., 2006, Martí-Renom et al., 2000, Sali and Blundell, 1993, Fiser et al., 2000) a test model was generated to verify the validity of the template and the alignment. This model was evaluated by computing its energy profile according to the DOPE-HR (high-resolution version of the Discrete Optimized Protein Energy) (Shen and Sali, 2006), smoothed via window averaging with a size of 15 residues. The profiles of template and model were compared (Supplementary Material S3), and further refinement was conducted in the region between Lys230 and Pro299, as well in all loop regions, using the slowest molecular-dynamics annealing available within MODELLER. For this step, 64 different base models were created and their secondary structure was refined independently 16 times. The resulting 1024 models were evaluated and ranked using DOPE-HR as well as the SOAP (Statistically Optimized Atomic Potentials) (Dong et al., 2013). The 10 best scoring models for each score were selected and evaluated based on the number of physical constraint violations present.

The best model of MurM_159_ was aligned with the previous MurM model (Fiser et al., 2003) or *W. viridescens* Femx homologues (Fonvielle et al., 2013, Biarrotte-Sorin et al., 2004) for visualisation and analysis in PyMOL (Version 2.1.0).

### Molecular docking of truncated Lipid II to MurM

A truncated Lipid II substrate (Supplementary Material: Figure S2) was created for initial molecular docking simulations. The truncated Lipid II was drawn in ChemDraw Professional (Version 17.1) and converted to a pdb file using Avogadro (Version 1.2.0). To prepare the ligand file for docking, the protonation state in H_2_O at pH 7.4 was computed. Subsequently the equilibrium geometry minimizing the potential energy was computed using the general amber force field (GAFF) (Wang et al., 2004) from within the Avogadro2 software (Hanwell et al., 2012). Molecular docking was conducted using AutoDock Vina (Trott and Olson, 2010), for which pdbqt files were generated from the pdb files of receptor model and ligands using AutoDock Tools (Morris et al., 2009). Initially the location of the binding site was verified by providing the algorithm with a search space that included the entire protein. Docking was then repeated by restricting the search space to the identified binding site, in order to obtain the final docked conformation.

### Coarse-grained molecular dynamics simulations

All coarse-grained simulations were carried out with the GROMACS package (Version 2018) and the Martini (Version 2.2) forcefield (Abraham et al., 2015, de Jong et al., 2012). Simulations at the coarse-grained and atomsitic resolutions were carried out at 313 K. For coarse-grained simulations, a stochastic velocity rescale thermostat with a coupling constant of 1.0 ps controlled the temperature.

The coordinates of the MurM homology model were used to generate a coarse-grained model using the ‘martinise.py’ script (de Jong et al., 2013). The protein was coarse grained to the ElNeDyn model (Periole et al., 2009) with an elastic network strength and cutoff of 500 kJmol^-1^nm^-2^ and 0.9 nm, respectively. The Lipid II model for inclusion in the membrane was parameterised using a united atom model (Gromos 53a6) generated by the Automated topology builder (ATB) web-interface. Following this, the coarse-grained mapping was decided iteratively and the bonded terms fitted with PyCGTOOL (Graham et al., 2017).

Since the pneumococcal membrane comprises a complex mixture of lipids, a simplified membrane composition was required for the simulations. In order to elucidate the effects of phosphatidylglycerol and cardiolipin on MurM, a non-pneumococcal lipid, phosphatidylethanolamine, was used as the majority lipid. Simulations were conducted with three different membrane systems (Supplementary Material: Table S1). System 1 was comprised of phosphatidylethanolamine and phosphatidylglycerol in a molar ratio of 75 % and 25 % respectively, system 2 contained phosphatidylethanolamine, phosphatidylglycerol and cardiolipin in a molar ratio of 72 %, 16 % and 12 % respectively and system 3 was comprised of phosphatidylethanolamine, phosphatidylglycerol and cardiolipin at a molar ratio of 72 %, 12 % and 16 % respectively. The membrane systems of size ~ 16 × 16 × 11.5 nm were generated with the CHARMM-GUI web interface (Jo et al., 2017). Each system was relaxed with a series of minimisation and equilibration steps with timesteps of 5 — 20 fs, for up to 30 ns. The equilibration steps utilised a semi-isotropic Berendsen barostat, with a 4.0 ps coupling constant (Berendsen et al., 1984). Following equilibration, Lipid II molecules (10 in total) were added to each membrane. The systems were then minimised and equilbriated (for 10 ns), followed by a 2 *μ*s production run to ensure sufficient mixing of all the lipid components. All production runs were carried out using a 10 fs timestep and a Parrinello-Rahman semi-isotropic barostat with a 12 ps coupling constant (Parrinello and Rahman, 1981). The Lennard-Jones potential was cutoff using the Potential shift Verlet scheme at long ranges. The reaction field method (Tironi et al., 1995) was used for electrostatics calculations, with dielectric constants of 15 and infinity for charge screening in the short- and long-range regimes, respectively. The short-range cutoff for non-bonded and electrostatic interactions was 1.2 nm. Once lipid mixing was ensured, the size of each system was increased to ~32 nm in the dimension perpendicular to the membrane and MurM was added in a random orientation around 8 nm above each membrane. Biologically relevant salt concentrations (0.15 M NaCl) were added and 10 % of the water molecules were changed to antifreeze particles to prevent localised freezing during simulations. After minimisation and 1 ns of equilibration, during which the protein backbone was restrained with 1000 kJmol^-1^nm^-2^ harmonic restraints, 6 × 5 μs production runs were generated per membrane composition (Supplementary Material: Table S1).

### All-atom molecular dynamics simulations

Atomistic simulations were conducted using the CHARMM36m forcefield (Huang et al., 2017). The Lipid II model used here was also used in previous work (Witzke et al., 2016), while all other lipid models were obtained from the CHARMM-GUI membrane builder module (Jo et al., 2008). For each coarse-grained membrane system, two repeats were chosen where: 1) the the last frame of the production run had a distinct orientation of MurM, relative to the membrane 2) MurM adhered to the membrane surface (Supplementary Data S2). The last frame of the chosen coarse-grained repeats were then backmapped to the all-atom model, using the backward script (Wassenaar et al., 2014). Unfavourable ring conformers were corrected by carrying our minimisation and equilibration steps with dihedral restraints of 25000 kJmol^-1^rad^-2^ on key ring torsions. After the transformation was carried out, each system was cropped in the z dimension to a height of 16.5 nm, to remove unnecessary H_2_O molecules.

Each system was minimised and equilibrated for a total of 1 ns, while the backbone of the protein was restrained with 1000 kJmol^-1^nm^-2^ harmonic restraints. Two production runs of 250 ns were carried out for each system. During the production runs a timestep of 2 fs was used, and the pressure (1 bar) regulated with a semi-isotropic Parrinello-Rahman barostat, with a coupling constant of 5.0 ps. The Lennard-Jones potential was cutoff with the Force-switch modifier from 1.0 to 1.2 nm. The short range cutoff for the electrostatic interaction was also 1.2 nm and the Particle mesh Ewald (PME) algorithm (Darden et al., 1993) was used for the long-range regime.

Analysis was carried out over the final 100 ns of each simulation, unless stated otherwise. All simulations were visualised using Visual Molecular Dynamics (VMD) or PyMOL (Version 2.2.0). Other analysis tools were written with a combination of GROMACS tools and in house scripts, that utilised the python module MDAnalysis (Gowers et al., 2016). The depletion/enrichment (D-E) indices were determined by first counting the number of lipids with a centre of geometry within 1.4 nm of the protein and then comparing this number to the number expected in the bulk of the membrane, using the procedure described by Corradi et al. (2018). The D-E index was obtained by dividing the lipid composition in the 1.4 nm shell around the protein by the bulk membrane composition. Thus a D-E index >1 indicates enrichment, while a D-E index <1 indicates depletion. The D-E index was determined for the last 100 ns of each simulation in 50 ns blocks for all repeats. For a given membrane composition, 8 D-E indicies were obtained for each lipid, from which the average and standard errors was calculated. The enrichment maps were generated by first determining the 2D density map of the membrane using the GROMACS tool densmap. Following this, the enrichment was determined using the procedure described by Corradi et al. (2018).

## Results

### X-ray crystallography and structure determination of S. aureus FemX

The crystal structure of *S. aureus* FemX was solved to a resolution of 1.62 Å and the structure was deposited in the PDB with accession number: 6SNR. A summary of the data collection and refinement statistics is given in Table 1.

**Table 1:** Summary of crystallographic data collection and refinement statistics from the *S. aureus* FemX structure. The highest resolution bin of data is indicated by square parentheses. Numbers in square parentheses refer to values in the highest resolution shell. ^*a*^*R*_sym_ = *S*_j_*S*_h_|*I*_h,j_ – < *I*_h_ > |*S*_j_*S*_h_ < *I*_h_ > where *I*_h,j_ is the is the jth observation of reflection h, and I_h_ is the mean intensity of that reflection. ^*b*^*R*_cryst_ = *S*||*F*_obs_| – |*F*_calc_||/*S*|*F*_obs_| where *F*_obs_ and *F*_calc_ are the observed and calculated structure factor amplitudes, respectively. ^*c*^*R*_free_ is equivalent to *R*_cryst_ for a 4% subset of reflections not used in the refinement (Brunger, 1992). ^d^DPI refers to the diffraction component precision index (Cruickshank, 1999). ^e^As calculated by Molprobity (Williams et al., 2018).

| Data collection | FemX |
| --- | --- |
| Synchrotron radiation, detector and wavelength (Å) | Pilatus 6M-F, 0.920 |
| Unit cell (a, b, c (Å), $\alpha$, $\beta$, $\gamma$ (°)) | 45.01, 83.62, 133.93, 90.0 |
| Space group | P2<sub>1</sub>2<sub>1</sub>2<sub>1</sub> |
| Resolution (Å) | 52.27-1.62 [1.66-1.62] |
| Observations | 422,822 [29,596] |
| Unique reflections | 65,058 [4,782] |
| I/s(I) | 15.7 [2.6] |
| R<sub>sym</sub><sup>a</sup> | 0.065 [0.567] |
| R<sub>meas</sub> | 0.078 [0.690] |
| R<sub>p.i.m</sub> | 0.031 [0.273] |
| Completeness (%) | 99.7 [99.8] |
| Refinement |  |
| Non-hydrogen atoms | 3,397 (including 177 waters) |
| R<sub>cryst</sub><sup>b</sup> | 0.221 [0.262] |
| Reflections used | 61,691 [4,531] |
| R<sub>free</sub><sup>c</sup> | 0.262 [0.296] |
| Reflections used | 3,294 [244] |
| R<sub>cryst</sub> (all data)<sup>b</sup> | 0.222 |
| Average temperature factor (Å<sup>2</sup>) | 26 |
| Rmsds from ideal values |  |
| Bonds (Å) | 0.013 |
| Angles (°) | 1.5 |
| DPI coordinate error (Å)<sup>d</sup> | 0.098 |
| Ramachandran Plot<sup>e</sup> |  |
| Favoured (%) | 98.0 |
| Outliers (%) | 0.0 |

The final solved structure of *S. aureus* FemX, contains two domains; a globular domain and a coiled-coil domain. Similarly to FemA (Benson et al., 2002), the globular domain can be divided into two subdomains. Each subdomain contains a central five-stranded mixed polarity β-sheet surrounded by four α-helices. Subdomain 1A comprises residues 1-145 and 384-421 whilst subdomain 1B comprises residues 146-234 and 298-383. Unfortunately, residues 403-421 were not present in the density. The coiled-coil domain consists of two antiparallel α-helices, comprising residues 235-297. FemX and FemA can be superimposed onto each other with a root-mean-square deviation (RMSD) of ~2.7 Å over 384 residues. Similarly to FemA, FemX has a deep L-shaped channel of about 20 x 40 Å located along side the globular domain and mainly in subdomain 1B. This channel comprises a peptidoglycan precursor binding site which was previously identified in *S. aureus* FemA (Benson et al., 2002). The identity of this peptidoglycan precursor binding site has been further confirmed by the crystallography studies of *W. viridescens* FemX complexed with substrates (Biarrotte-Sorin et al., 2004).

### Homology modelling of S. pneumoniae MurM

The structures of two MurM homologues, *S. aureus* FemA and *W. viridescens* FemX were solved previously by X-ray crystallography (Benson et al., 2002, Fonvielle et al., 2013, Biarrotte-Sorin et al., 2004). The *S. aureus* FemA structure was subsequently used as a template for homology modelling of MurM by Fiser et al. (2003). However, *S. aureus* FemX possesses a higher sequence identity to MurM (Table 2), and is also functionally more homologous to MurM. *S. aureus* FemA appends the second and third amino acid residues of the cross-bridge to the α-amino group of a glycyl residue appended to the e-amino group of the stem peptide L-lysyl residue of the Lipid II precursor. In contrast, *S. aureus* FemX, similarly to *S. pneumoniae* MurM, appends the first amino acid of the cross-bridge to the Lipid II precursor (Matsuhashi et al., 1967, Schneider et al., 2004). Therefore, given the difficulties in obtaining MurM crystals, we were motivated to solve the structure of its functional homologue (*S. aureus* FemX) by X-ray crystallography, to access by *in silico* approaches the structure of MurM.

**Table 2:** Sequence identity and similarity of *S. pneumoniae* MurM and three homologues. *S. pneumoniae* MurM sequence was aligned with three homologues *S. aureus* FemA, *S. aureus* FemX and *W. viridescens* FemX using EMBOSS Needle (Madeira et al., 2019).

| Homologue | Sequence Identity (%) | Sequence Similarity (%) |
| --- | --- | --- |
| *S. aureus* FemA | 21.6 | 40.3 |
| *S. aureus* FemX | 27.3 | 44.3 |
| *W. viridescens* FemX | 23.1 | 27.5 |

Using the newly solved structure of *S. aureus* FemX we generated a new homology model for MurM which consists of a globular domain comprising two subdomains and a coiled-coil helical arm (Figure 2. Each subdomain comprises two twisted *β*-sheet cores surrounded by alpha helices; subdomain 1A is formed of residues 1-153 and 382-401, whilst subdomain 1B is made up of residues 154-241 and 294-381. The coiled-coil domain comprises residues 242-293. Whilst the new MurM homology model is similar to the previous model (Fiser et al., 2003), the RMSD of the two models is ~3.8 Å over 368 residues, indicating that there are some key structural differences, namely; loss of N-terminal β 1, and antiparallel β6/β 13 from the previous model; addition of α5 and β11/β 12; and presence of *α* helical secondary structure at the C-terminal end of the new MurM model.

**Figure 2:**
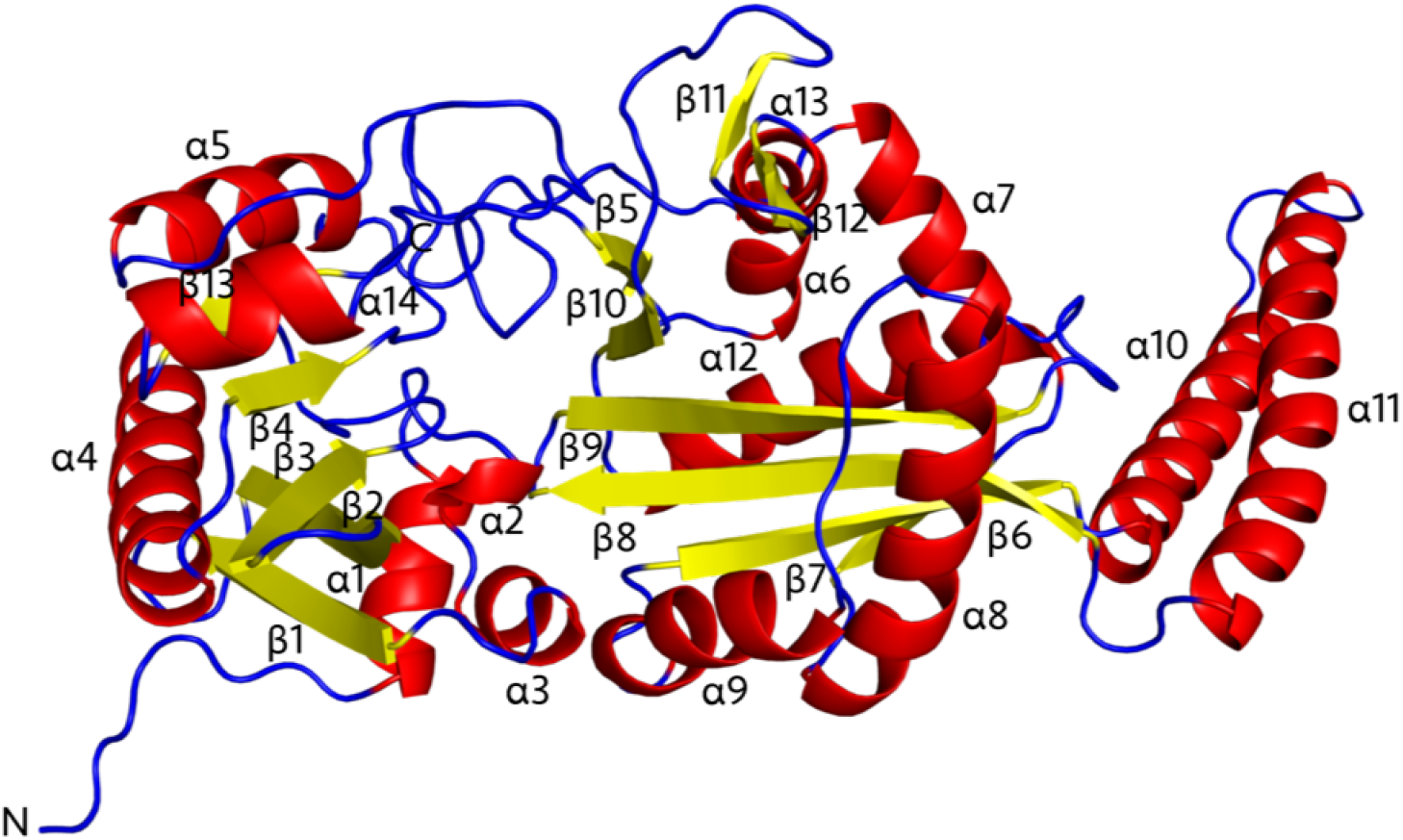
Cartoon representation of MurM predicted structure. 14 *α*-helices (red), 12 *β*-sheet (yellow) and unstructured regions (blue). Best model obtained based on SOAP and DOPE scores following homology modelling using MODELLER with *S. aureus* FemX as a template.

### Identification of a possible Lipid II binding site of MurM

The new MurM model revealed a binding pocket which was not present in the previous model of MurM. Structural comparison between *W. viridescens* FemX co-crystallised with its UDP-Mur*N*Ac-pentapeptide substrate and the new MurM model, allowed identification of a Lipid II binding site that corresponds with those identified previously in *S. aureus* FemA and FemX, as well as *W. viridescens* FemX. When the new MurM model and the *W. viridescens* FemX were aligned and overlaid, the newly identified MurM binding site appeared to easily accommodate the soluble UDP-Mur*N*Ac-pentapeptide substrate well (Figure 3a). The following 8 residues; Tyr103, Lys36, Asn38, Trp39, Thr209, Arg211, Try215 and Tyr256, were independently proposed to be involved in substrate binding in both *W. viridescens* FemX structures (Biarrotte-Sorin et al., 2004, Fonvielle et al., 2013). The corresponding MurM residues, defined as having residues which have similar properties, and occupying a similar location and orientation in physical space, with side chains facing the binding pocket, were identified in the MurM structure as Phe103, Lys35, Trp38, Arg215 and Tyr219, therefore these residues may also be important for substrate binding in MurM.

**Figure 3:**
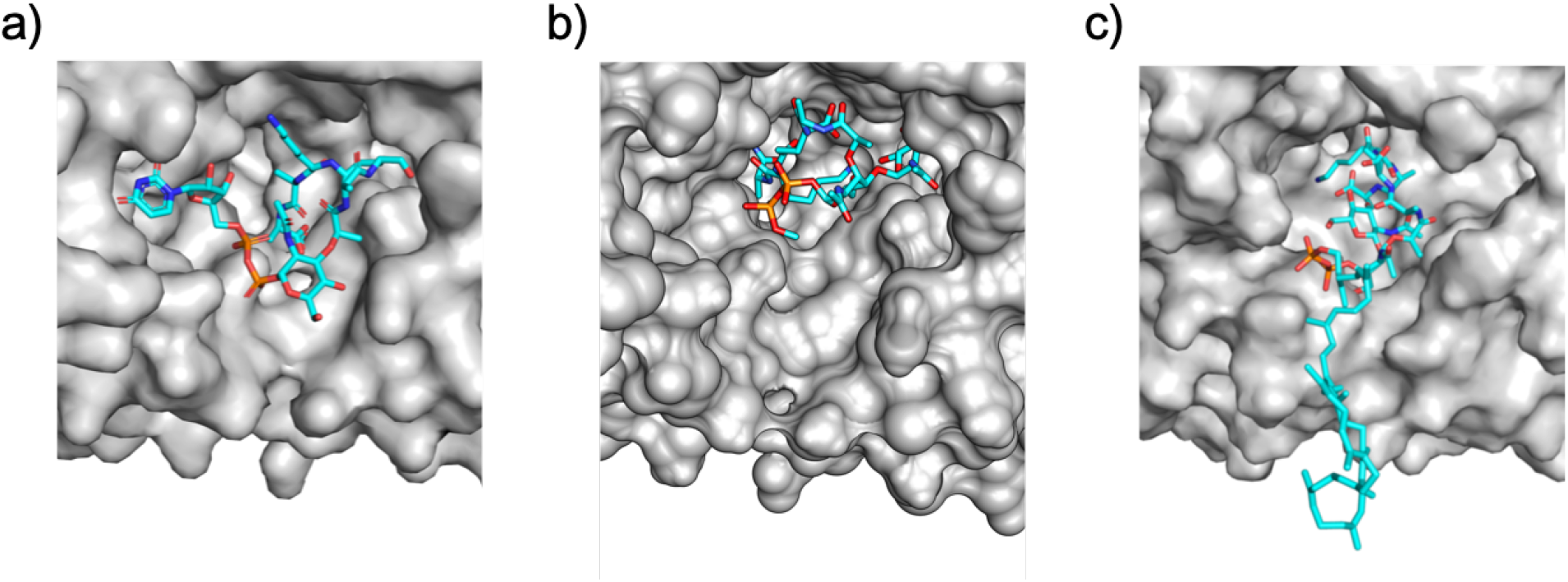
Surface representation of MurM binding site. a) MurM 159 model aligned and overlaid with the UDP-Mur*N*Ac-pentapeptide substrate which was co-crystallised with *W. viridescens* FemX b) MurM_159_ model with truncated Lipid II docked in the binding site, using AutoDock Vina c) MurM_159_ model with Lipid II in the binding site, from membrane simulations. Figures were created with PyMOL (Version 2.2.0) and Chimera (Version 1.13.1).

Next molecular docking using AutoDock Vina (Trott and Olson, 2010) was conducted to independently investigate docking of the Lipid II substrate to the new MurM model. Lipid II is a large molecule that is in general unsuitable for molecular docking studies. In addition, the lipid tail is embedded in the membrane, and so is not itself available for binding to MurM. Therefore, a truncated Lipid II molecule, comprised of a methyl capped diphosphate Glc*N*Ac-Mur*N*Ac-pentapeptide, was used for these docking experiments (Supplementary Material: Figure S2).

When AutoDock Vina was allowed to search the entire protein surface of MurM, all docking results returned were within the identified binding site, indicating that there are no other suitable binding sites on the protein. The search was then restricted to the binding site and the top ten results were obtained. The top five results obtained all had identical identical binding affinities of −7.3 kcal.mol^-1^. Two docking orientations, whereby the phosphates are located deep within the binding pocket, would be physically impossible for the natural substrate (Lipid II) *in vivo*, since the membrane-embedded prenyl lipid tail is appended via the phosphate. The remaining 3 docking orientations, all orient the phosphates close to the opening of the binding site with the pentapeptide chain disappearing deep into the binding pocket. The exact orientation of the pentapeptide chain is variable, indicating that the binding site is spacious and that Lipid II may be accommodated in a number of different possible orientations. Figure 3b) shows one conformation in which the docking of truncated Lipid II is similar to the orientation of the soluble UDP-Mur*N*Ac-pentapeptide from *W. viridescens* FemX overlaid with MurM (Figure 3a), the remaining four substrate orientations with binding affinities of −7.3 kcal.mol^-1^ are shown in the Supplementary Material (Figure S5). A key limitation of docking is that it considers the protein as rigid; therefore, multiple substrate orientations may indicate that conformational changes within the binding site may occur upon substrate binding or during catalysis.

### Interactions between MurM and the lipid bilayer using Molecular dynamics

Molecular dynamics simulations were used to model the interactions of MurM with the lipid bilayer. Coarse grain simulations were conducted six times for each of three membrane systems. In most runs MurM readily associated with the membrane in <3 μs and remained unchanged for the remaining 2 μs (Supplementary Material: Figure S4). In addition, Lipid II in the same leaflet as the peripheral MurM was found to cluster around the protein; more than 50 % of Lipid II molecules were located within 2 nm of the MurM.

MurM associated with the membrane in a number of different orientations. However, in 1 out of 18 unbiased simulations, the MurM was oriented such that the Lipid II molecule was located in the putative binding site, which demonstrates that the Lipid II is able to successfully enter this binding site even on the short timescale of a MD simulation. Back mapping of this simulation to all-atom resolution allowed the binding site to be explored in more detail. The overall orientation of the Lipid II in the binding site is very similar to the docking of the truncated substrate (Figure 3a).

Similarly to the molecular docking findings, this MD simulation shows that the MurM binding site is flexible and allows the Lipid II molecule to adopt a wide variety of conformations (Figure 4). This may suggest that binding of a second substrate or a large conformational change may be required for catalysis.

**Figure 4:**
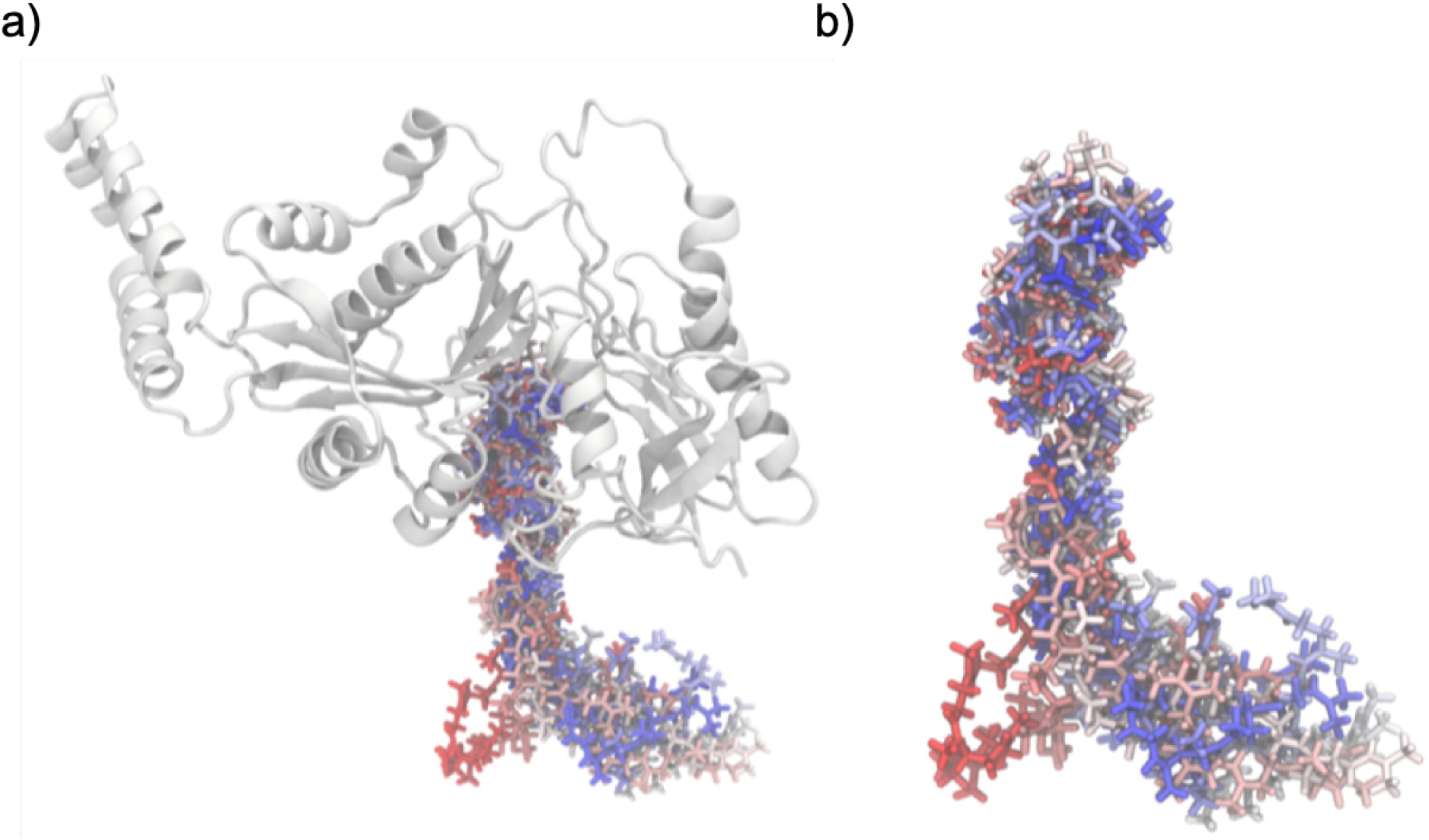
Different conformations of Lipid II inside MurM binding site. Lipid II binding in system 5 coloured on a BWR scale with respect to simulation time.

### Interactions between MurM and membrane phospholipids

Molecular dynamics simulations were used to investigate the effects of membrane phospholipids (cardiolipin and phosphatidylglycerol) on MurM at the cytoplasmic membrane interface. Figure 5 shows that upon association of MurM with the cytoplasmic membrane, there was no effect on the distribution of phosphatidylglycerol or phosphatidylethanolamine. However, in membranes containing 8 % or 12 % cardiolipin, cardiolipin was enriched at the MurM:membrane interface, whilst the phosphatidylethanolamine and phosphatidylglycerol distributions remained largely unaffected. The importance of these observations were considered *in vitro* by measuring the enzymatic activity of MurM in the presence of varying concentrations of cardiolipin or phosphatidylglycerol. These enzymatic studies show that cardiolipin activates MurM, whilst phosphatidylglycerol inhibits MurM in a concentration dependent manner. Figure 5e shows the enzymatic activation of MurM with respect to cardiolipin concentration, a 9.1-fold activation of MurM was achieved, with 50 % activation occurring at 0.4 mM cardiolipin. Figure 5f shows that the activity of MurM could be completely inhibited by phosphatidylglycerol, with an IC_50_ of 0.2 mM. Furthermore, Hill coefficients of 2.7±0.3 and 2.8±0.2 for cardiolipin and phosphatidylglycerol respectively, indicate that both these phospholipids exhibit their effects on MurM in a cooperative manner. Phosphatidylethanolamine, used in the construction of the model pneumococcal membrane to which MurM bound, when tested at a concentration of 0.72 mM, only slightly activated MurM activity by 0.32-fold (duplicate determination with a difference of ¡10 %. In comparison, 0.72 mM cardiolipin activated MurM by 8-fold (Figure 5e). Therefore, the impact of phosphatidylethanolamine on the disposition of MurM relative to its interaction with Lipid II and the phospholipid bilayer could be neglected.

**Figure 5:**
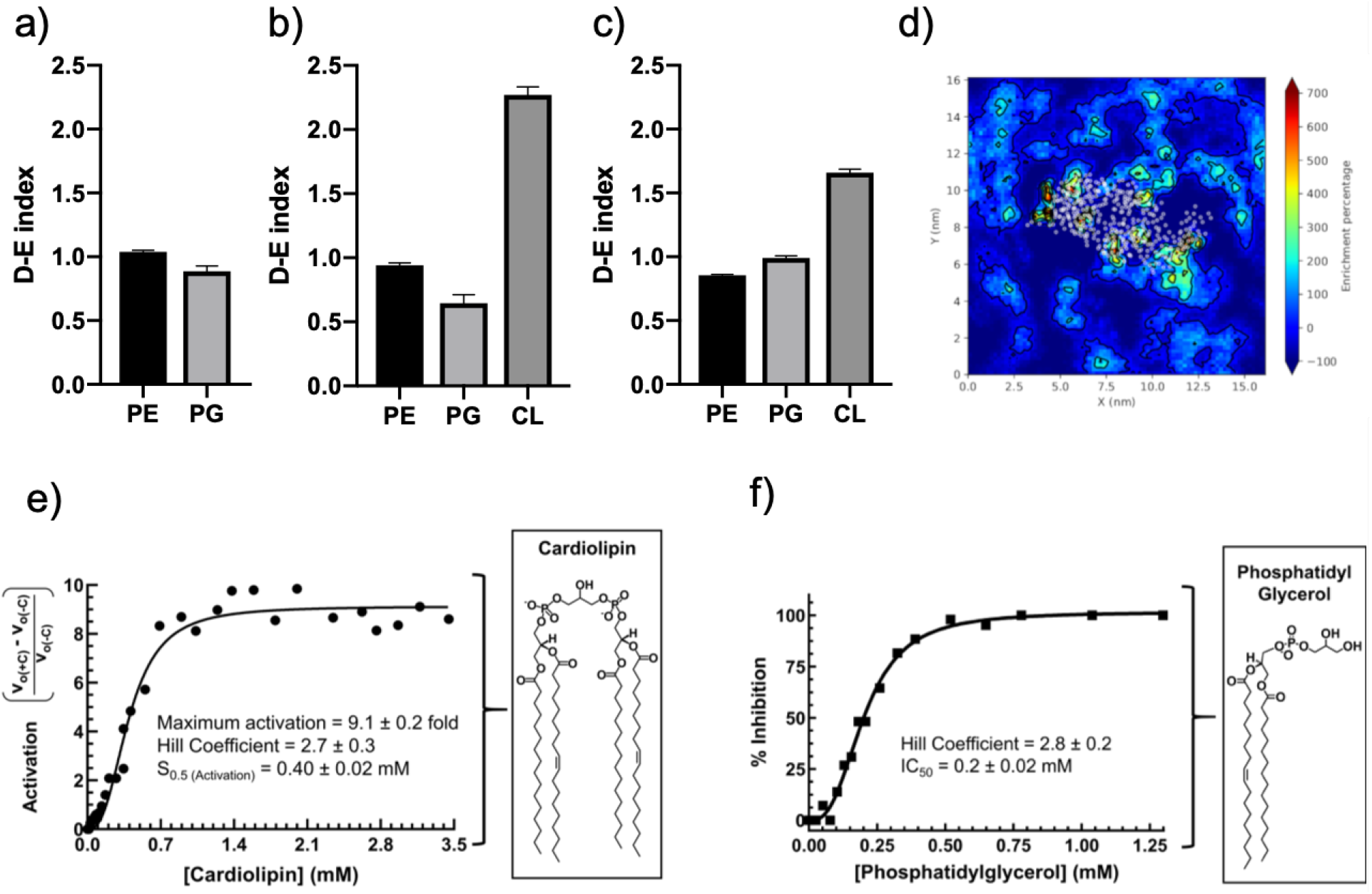
Interactions between MurM and membrane phospholipids. Depletion-enrichment (D-E) indices for phosphatidylethanolamine (PE), phosphatidylglycerol (PhG) and cardiolipin (CL) occurring within a 1.1 nm perimeter of the MurM protein for a) Systems 4-5 (molar ratio of 75% phosphatidylethanolamine and 25% phosphatidylglycerol), b) System 6-7 (molar ratio of 76% phosphatidylethanolamine, 16% phosphatidylglycerol and 8% cardiolipin) and c) System 8-9 (molar ratio of 72% phosphatidylethanolamine, 12% phosphatidylglycerol and 16% cardiolipin). The D-E index was determined from 150-250 ns in 50 ns blocks for all repeats for a total of 8 values per plot. d) Example of a depletion-enrichment map with MurM at the membrane. White dots represent the center of geometry of each protein residue, and the percentage enrichment of phospholipid is indicated by the colour. e) Activation of MurM was calculated as the product of subtraction of MurM velocity in the absence of cardiolipin (v_o(-c)_) from MurM velocity in the presence of cardiolipin (v_0(+c)_) divided by v_0(-c)_ and was plotted versus cardiolipin concentration. f) Inhibition of MurM was calculated as ((v_0(-PG)_) – (v_0(+PG)_))/v_0(-PG)_ x 100 (where PG denotes phosphatidylglycerol) and was plotted versus phosphatidylglycerol concentration. Data were fitted as described in the text. GraphPad Prism (Version 7.0c) and Matplotlib (Version 3.0.3) were used for data analysis and figure preparation.

## Discussion

The crystal structure of *S. aureus* FemX has allowed the us to generate an improved homology model of MurM leading to the identification of a putative Lipid II binding site. Fiser et al. (2003) proposed a different MurM model and speculated about an alternative binding site based upon structural and functional analogy between MurM and N-myristoyltransferase (NMT) proteins. However, whilst the substrates of both NMT proteins and MurM are lipids, they are contextually very different. The NMT proteins are cytoplasmic proteins that contain a deep, narrow pocket which is highly specific for the myristoyl fatty acyl chain (Wright et al., 2010, Heuckeroth et al., 1988). In contrast, MurM binds the disaccharide head group and pentapeptide side chain of Lipid II, and the undecaprenyl C55 lipid tail is embedded in the membrane. Despite similarities with NMT proteins, our newly identified substrate binding site more closely resembles those of *W. viridescens* FemX, *S. aureus* FemX and FemA.

The orientations of truncated Lipid II in docking studies and Lipid II in molecular dynamics simulations are strikingly similar to each other and also the orientation of UDP-Mur*N*Ac-pentapeptide substrate co-crystallised in *W. viridescens* FemX. This is consistent with the observation that, although inefficient compared to Lipid II, UDP-Mur*N*Ac pentapeptide is a MurM substrate (Lloyd et al., 2008). In all cases, the diphosphates are near the surface of the protein, and the protruding pentapeptide reaches into the binding pocket with the third-position lysine on the left-hand side of the binding pocket. In addition, previous studies suggest that the height of the Lipid II head group is 19 Å (Ganchev et al., 2006), and the binding pocket of this model was measured to be 15 Å. Since the Lipid II head group is flexible, and the binding site provides enough room for the substrate to bend, these measurements are consistent. Together with our findings, this strongly supports the identification of this newly identified cavity as the Lipid II binding site and suggests that the Lipid II binds to MurM in an orientation similar to that of *W. viridescens* FemX binding to its substrate.

Alanylphosphatidylglycerol synthase (PDB ID 4v34) similarly to MurM, also utilises both lipid and alanyl-tRNA^Ala^ substrates (Hebecker et al., 2015), in order to successfully bring these two substrates together for catalysis, it possesses two binding sites, located on opposite sides of the protein which are connected by a channel. The protein itself provides a barrier between the hydophobic lipid and the hydrophillic tRNA, such that they do not come into close proximity with each other. The negatively charged surface patch identified previously (Fiser et al., 2003) remains present on this new homology model of MurM, and is located on the opposite side of the protein with respect to the Lipid II binding site, which is located within a positively charged surface patch (Supplementary Material: Figure S6). This negative patch is unsuitable for the binding of negatively charged tRNA and so it is unlikely that MurM shares the same mechanism of action as alanylphosphatidylglycerol synthase. The negatively charged surface patch may however be important for protein:protein interactions occurring either at the cell surface or in the cytoplasm.

These modelling studies reveal that the stem peptide protrudes perpendicular to the surface of the membrane into the active site of MurM. Therefore, in order for alanyl-tRNA^Ala^ to simultaneously interact with MurM, whilst it is located over its lipid substrate, the highly negatively charged hydrophilic tRNA would have to be brought into close proximity with the negatively charged phospholipid head groups and/or the hydrophobic phospholipid tails below them. Given that this would be a highly unfavourable interaction, we propose an alternative ‘ping-pong’ mechanism of action for MurM whereby MurM is initially aminoacylated by alanyl-or seryl-tRNA in the cytoplasm before travelling to the cell membrane for transfer to Lipid II. Whilst the MurM is in the cytoplasm and not interacting with the membrane, the positively charged patch, located at the newly proposed Lipid II binding site, may facilitate interaction with a polyanionic substrate such as tRNA. Once this has occurred, subsequent interaction of the aminoacyl-MurM with the surface of the membrane could accommodate the correct and catalytically productive interaction of aminoacylated-MurM with lipid II. Although this proposed mechanism is at variance with the sequential mechanism of catalysis proposed for *W. viridescens* FemX (Hegde and Blanchard, 2003), in this case, both substrates were highly hydrophilic nucleotide or polynucleotide derivatives in the same cellular sub-compartment and are therefore without biophysical impediment with regard to their proximity during catalysis. Here, with regard to MurM, the chemical properties and location of both substrates indicate an advantage to a mechanism which avoids their simultaneous binding.

The pneumococcal peptidoglycan is heterogeneous with respect to its composition of directly and indirectly cross linked stem peptides. It remains unclear as to whether the activity of MurM, and therefore the generation of indirect cross-links is distributed equally around the entire cell surface, or whether it is localised to specific sites. Phospholipids are known to be involved in the spatial and temporal biochemistry of cells (Lin et al., 2019), and cardiolipin was shown to be enriched at the poles and septa of *E. coli* and *Bacillus subtillis*, localising specific membrane-associated proteins to these regions (Bramkamp and Lopez, 2015). Our simulations indicate that, whilst cardiolipin enrichment occurs within the membrane in the presence of MurM, this phospholipid is not essential for membrane association of MurM to occur. Therefore, it remains uncertain as to whether *in vivo* cardiolipin is highly concentrated in patches in the membrane and is used to recruit MurM to that location, or whether association of MurM with the membrane drives the enrichment of cardiolipin in the membrane.

Despite this uncertainty, we show that cardiolipin stimulates the enzymatic activity of MurM, and whilst it is not clear if this increased activity is as a result of a direct effect on the protein or the Lipid II substrate, or both, the spatial association of cardiolipin to the MurM protein suggests that at least some of this effect may be due to direct interactions with the MurM protein. Cardiolipin has previously been found to bind to and activate a wide range of proteins including MurG (Boots et al., 2003), rat liver protein kinase N (Morrice et al., 1994, Peng et al., 1996), porcine heart AMP deaminase (Purzycka-Preis and Zydowo, 1987), rat liver multi-catalytic proteinase (Ruiz de Mena et al., 1993), *E. coli* glycerol-3-phosphate acyltransferase (Scheideler and Bell, 1989), *E. coli* dnaA (Sekimizu and Kornberg, 1988) and streptococcal hyaluronan synthases (Tlapak-Simmons et al., 1999a,b, 2004, Weigel et al., 2006, Tlapak-Simmons et al., 1998). This further supports the contention that cardiolipin affects MurM activity by directly interacting with MurM. Similar cardiolipin-mediated sigmoidal stimulatory effects have been seen with other streptococcal membrane proteins such as the hyaluronan synthases from *Streptococcus pyogenes* and *Streptococcus equismilis* (Tlapak-Simmons et al., 1999a,b, 2004, Weigel et al., 2006). In these examples, up to sixteen cardiolipin molecules are believed to associate with single hyaluronan synthase molecule (Tlapak-Simmons et al., 1998).

We also show that phosphatidylglycerol inhibits the catalytic activity of MurM, and that the concentration of this lipid in the membrane environment surrounding the MurM changes very little. Therefore, the inhibitory effect of phosphatidylglycerol may be exerted by altering the presentation of the Lipid II substrate to MurM, rather than by having a direct effect on the protein itself. It is possible that in *S. pneumoniae*, as in *E. coli* and *B. subtillis*, cardiolipin gathers in specific regions of the membrane, where it localises and up-regulates the activity of MurM, resulting in higher levels of indirect cross-linking in these regions.

Whilst MurM alone is not sufficient for penicillin resistance, the enzyme is crucial together with mosaic *S. pneumoniae* PBPs for the generation of a highly resistant phenotype. Deletion of *murM* from resistant strains resulted in a virtual abolition of penicillin resistance that could not be restored by *PBP* DNA. Indeed, additional *murM* DNA from a resistant strain was required for full expression of donor level penicillin resistance (Filipe and Tomasz, 2000, Smith and Klugman, 2001). Given the importance of MurM for penicillin resistance, the enrichment of cardiolipin at the MurM:membrane interface which activates MurM, and the inhibition of MurM activity by phosphatidylglycerol, may regulate the penicillin resistance phenotype imparted by MurM activity which may therefore be regulated by cardiolipin synthase activity. These findings have therefore revealed a crucial and hitherto unexplored area with regards to establishing a more complete understanding of penicillin resistance mechanisms and the influence on them by other areas of pneumococcal metabolism.

This new MurM structural model allowed identification of the Lipid II binding site and the contextual presentation of this substrate to MurM, characterised the impact of membrane phospholipids on MurM at the MurM:membrane interface, and may have spatial mechanistic implications for the catalytic activity of this protein. Molecular dynamics enabled the *in silico* investigation into MurM:membrane interactions, which are often overlooked when studying enzymes which act at the cytoplasmic membrane interface. The subsequent *in vitro* experiments on the importance of phospholipids for MurM activity, corroborate the *in silico* findings, supporting the role of phospholipids as an important contributor to the regulation of MurM at the membrane. These studies provide new insights into the structure of MurM which may guide future mutational studies, and allow a more detailed analysis of the structure-function relationship of this protein. This research contributes important findings towards achieving a more complete understanding of its role in pneumococcal penicillin resistance mechanisms.

## Acknowledgements

This work was supported in part by the Midlands Integrative Biosciences Training Partnership (MIBTP) BBSRC grant BB/J014532/1, and the Centre for Doctoral Training in Theory and Modelling in Chemical Sciences (TMCS DTC) EPSRC grant EP/L015722/1, as well as MRC grants G1100127, G0400848, MR/N002679/1 and BBSRC grant BB/N003241/1. The authors would like to acknowledge the help of the Media Preparation Facility in the School of Life Sciences at the University of Warwick. We would also like to thank Dr Allister Crow for his help and support with using PyMOL.

## Author Contributions

A.Y, A.J.L and D.I.R conceptualised this study, K.F constructed plasmids and purified proteins, K.J.H and V.F prepared samples and performed X-ray crystallography, V.F deposited the structure in the PDB. C.I.G and A.Y conducted homology modelling and docking studies, J.S conducted molecular dynamics studies, A.J.L cloned and purified proteins and conducted activity assays. A.Y conducted analysis of data and wrote the manuscript with contributions and manuscript reviews from J.S, C.I.G. C.G.D, D.I.R, A.J.L and S.K.

## Supplementary Material

**Figure S1:**
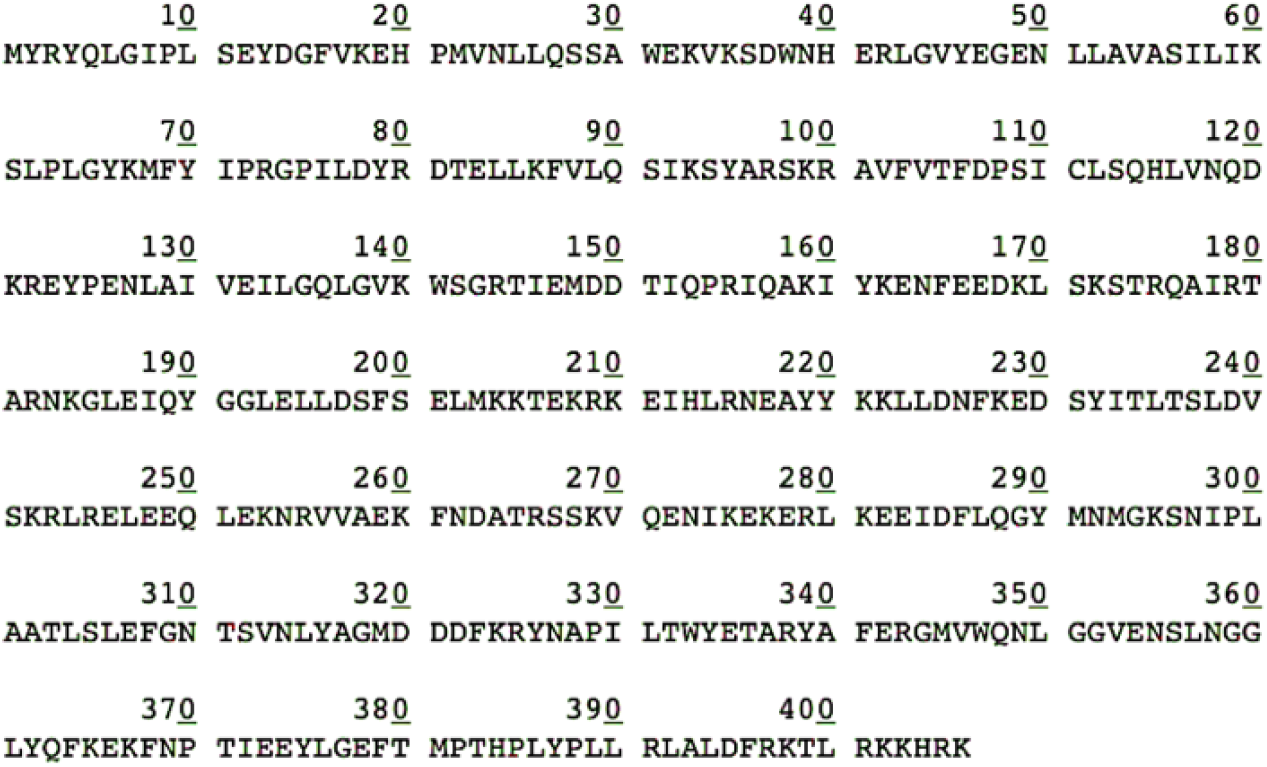
MurM159 protein sequence.

**Table S1:** Summary of course grained simulations.

| System | Membrane size (nm) | Time (μs) | Membrane % Composition | | |
| --- | --- | --- | --- | --- | --- |
|  |  |  | PE | PhG | CL |
| 1 | 16.1x16.1 | 6x5 | 75 | 25 | 0 |
| 2 | 16.1x16.1 | 6x5 | 76 | 16 | 8 |
| 3 | 16.3x16.3 | 6x5 | 72 | 12 | 16 |

**Table S2:** Summary of atomistic simulations. NB. CG label refers to the coarse grained system (Table S1) from which the simulation was constructed.

| System Label | Time (μs) | CG Label |
| --- | --- | --- |
| 4 | 2x0.25 | 1 r1 |
| 5 | 2x0.25 | 1 r2 |
| 6 | 2x0.25 | 2 r1 |
| 7 | 2x0.25 | 2 r6 |
| 8 | 2x0.25 | 3 r1 |
| 9 | 2x0.25 | 3 r4 |

**Figure S2:**
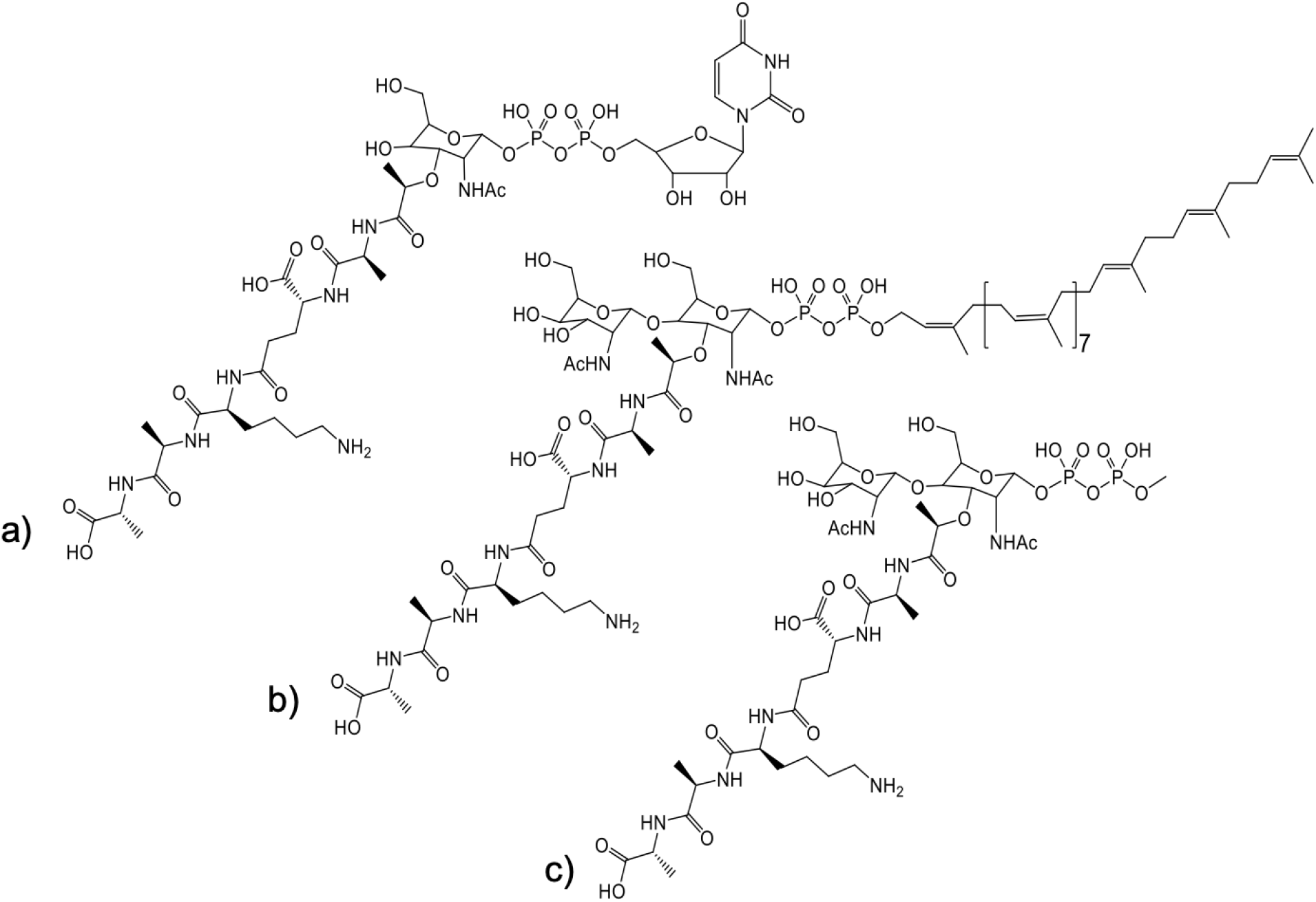
Structures of UDP-Mur*N*Ac-pentapeptide, Lipid II and the truncated Lipid II used for molecular docking studies. a) UDP-Mur*N*Ac-pentapeptide (Lysine variant) b) Lipid II c) Truncated Lipid II structure where the C55 prenyl chain has been replaced with a methyl group.

**Figure S3:**
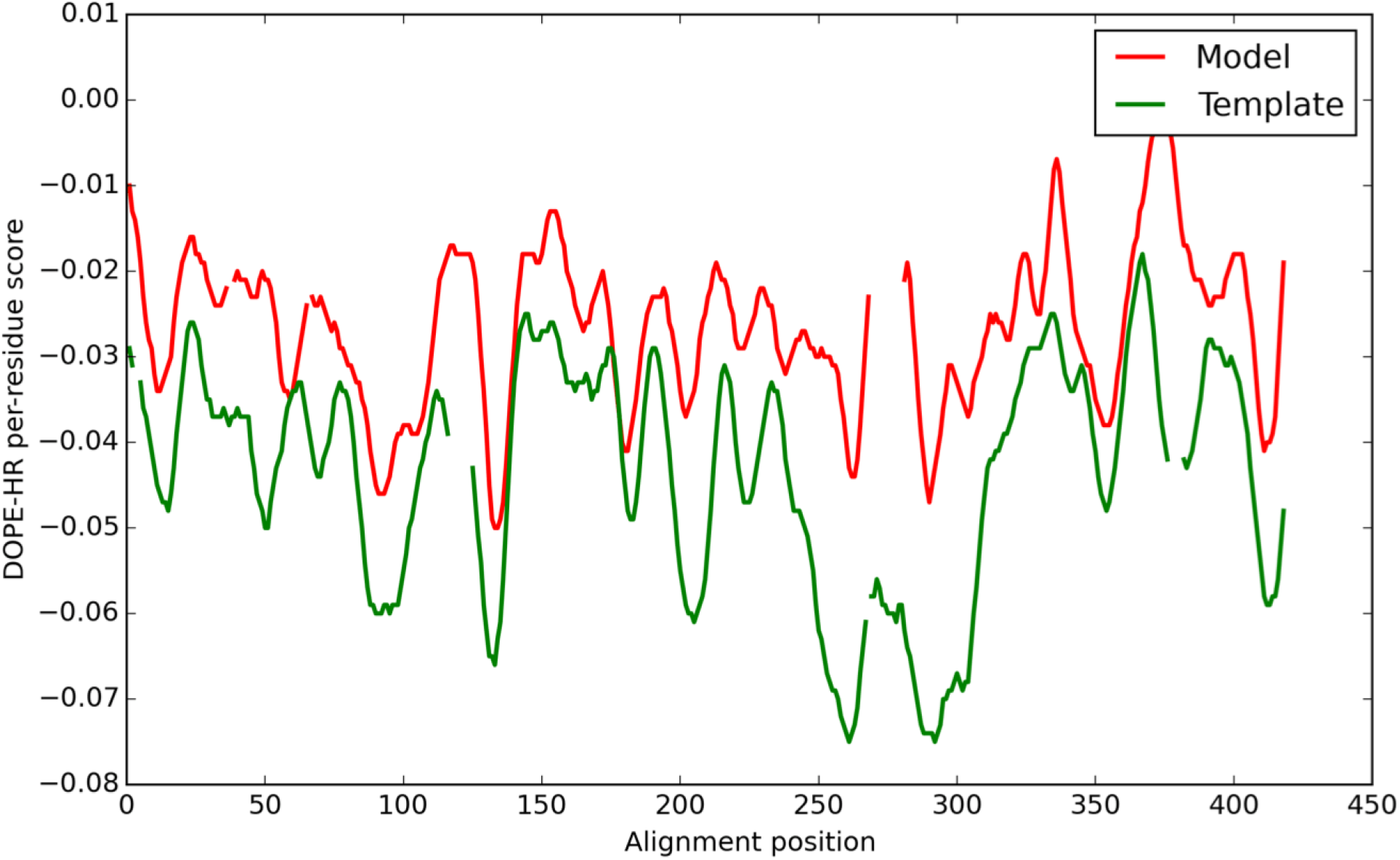
Discrete Optimized Protein Energy Profile for MurM and FemX. Comparison of DOPE-HR profiles for MurM model (red) and FemX template (green).

**Figure S4:**
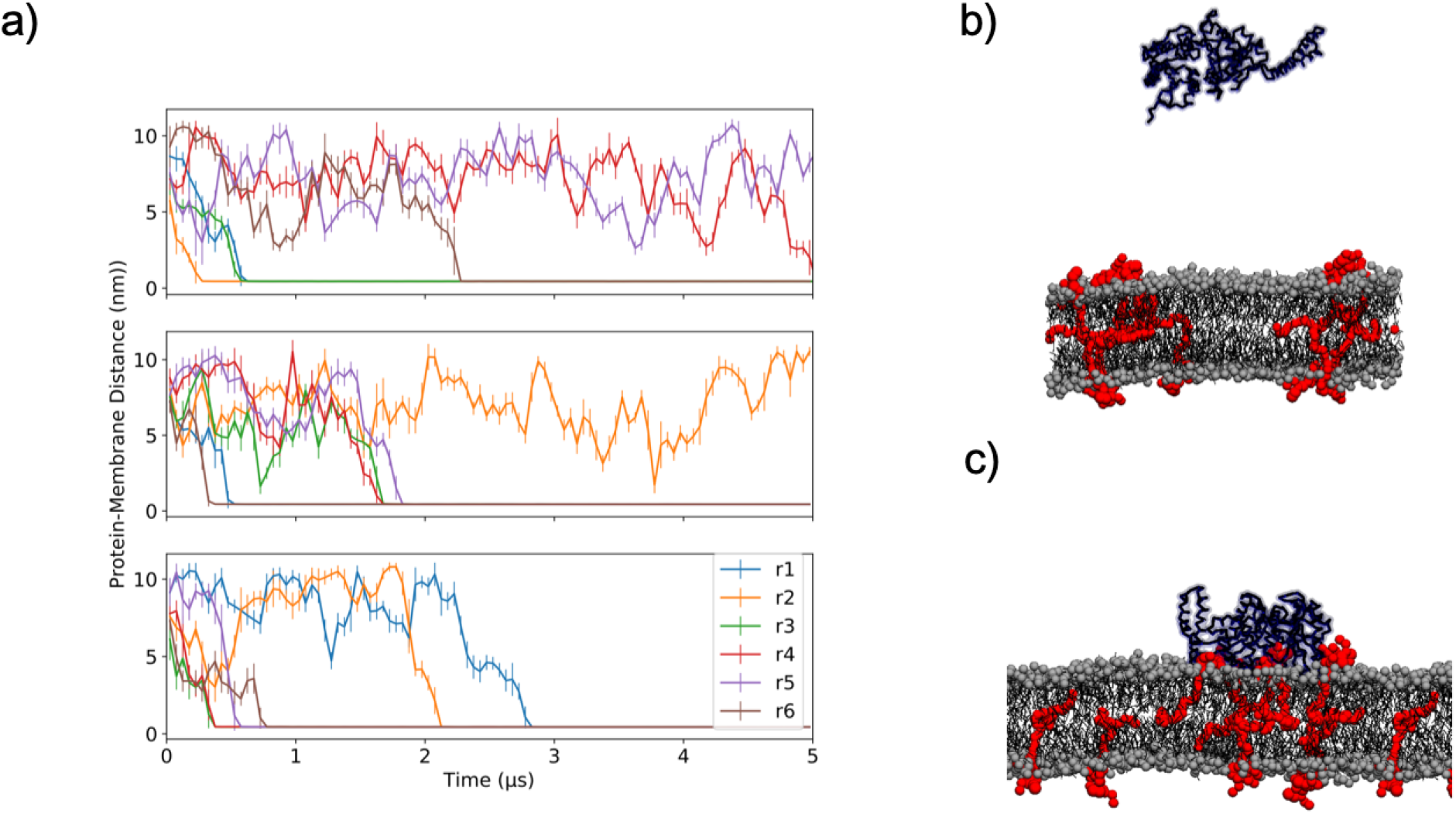
The association of coarse-grained MurM to the surface of the membrane. a) The minimum distance between MurM and the membrane surface for System 1 (top), System 2 (middle) and System 3 (bottom). Snapshots taken of the b) first and c) last frame of repeat 2 (r2), for System 1 (S1). Colour key: red = lipid II,blue = protein, and gray = membrane.

**Figure S5:**
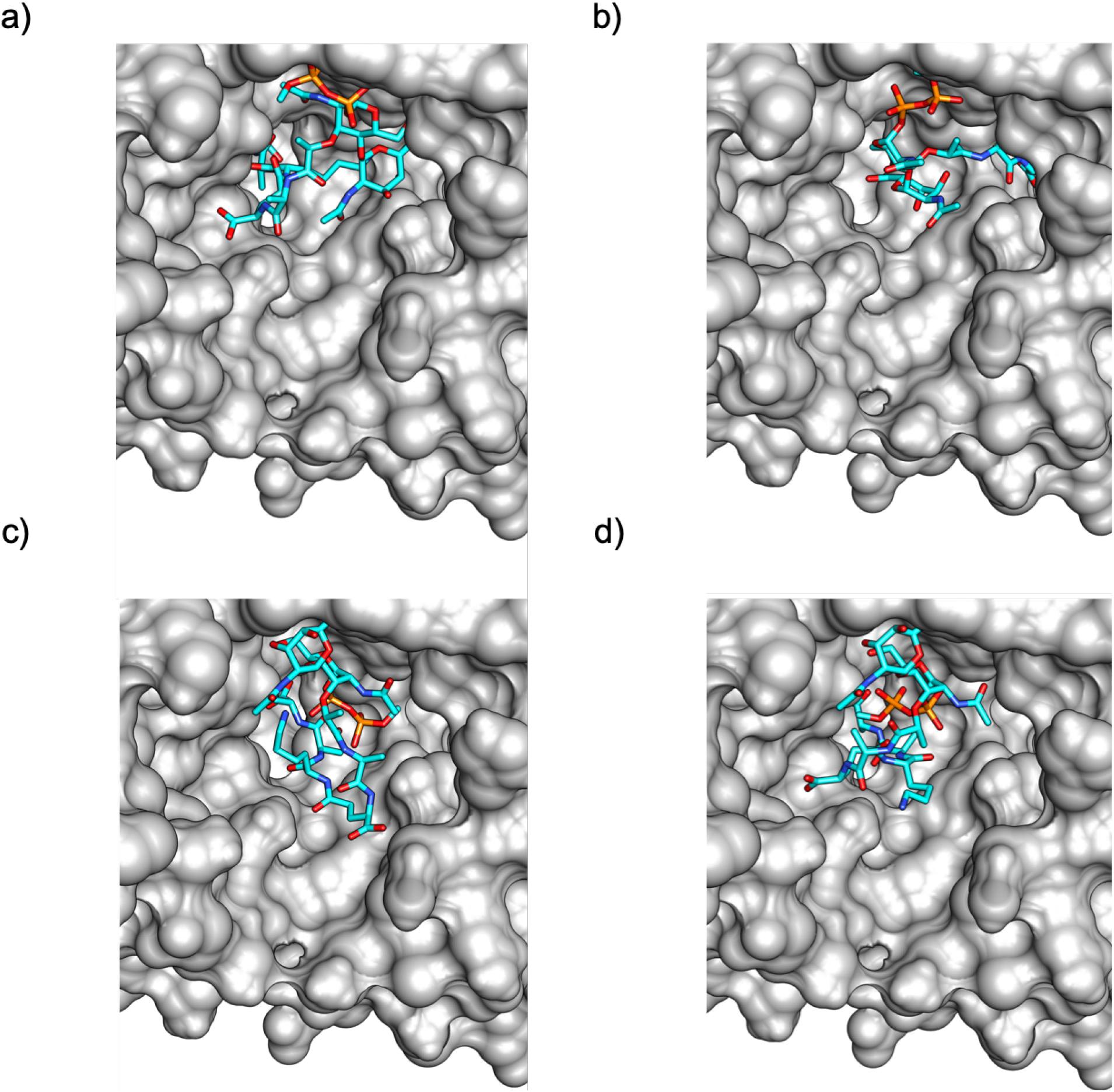
The remaining four highest scoring poses from molecular docking of truncated Lipid II to MurM. All possessed identical binding affinities of −7.3 kcal.mol^-1^. a) and b) show the phosphate group located near the entrance of the cavity, with the pentapeptide located deeper into the pocket. c) and d) show orientations that are not considered possible, since the phosphate group would be linked to the membrane embedded Lipid II, and this would prevent the phosphate from being located deep in the binding site as shown.

**Figure S6:**
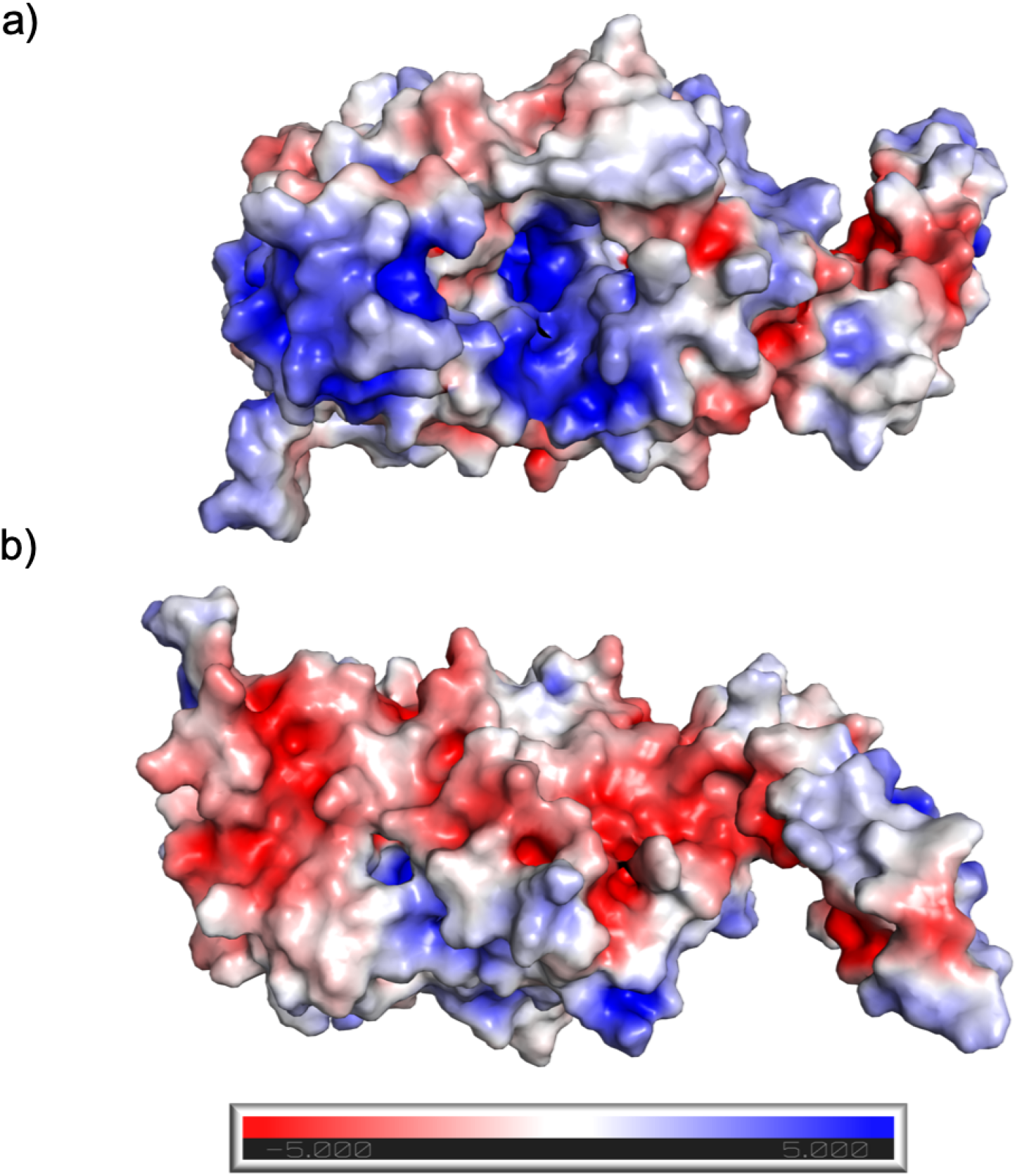
Electrostatic surface representation of MurM. a) MurM showing the proposed Lipid II binding site to be positively charged (Blue) b) MurM rotated 180, showing a negatively charged surface patch. Figure prepared in PyMOL (Version 2.2.0) using the APBS Electrostatics Pluggin.

